# Utilizing protein structure graph embeddings to predict the pathogenicity of missense variants

**DOI:** 10.1101/2024.11.15.623748

**Authors:** Martin Danner, Matthias Begemann, Miriam Elbracht, Ingo Kurth, Jeremias Krause

## Abstract

**Background:** Genetic variants can impact the structure of the corresponding protein, which can have detrimental effects on protein function. While the effect of protein truncating variants is often easier to evaluate, most genetic variants are missense variants. These variants are mostly single nucleotide variants which result in the exchange of a single amino acid. The effect on protein function of these variants can be challenging to deduce. To aid the interpretation of missense variants a variety of bioinformatic algorithms have been developed, yet current algorithms rarely directly use the protein structure as a feature to consider.

**Results:** We developed a machine learning workflow that utilizes the protein-language-model ESMFold to predict the protein structure of missense variants, which is subsequently embedded using graph autoencoders. The generated embeddings are used in a classifier model which predicts pathogenicity. We provide evidence that the generated graph embeddings improve classification accuracy of a XGBoost pathogenicity predictor, which should lead to a wide applicability for human genetic diseases. Additionally, we explored different levels of abstraction of the graph embeddings and their influence on the classifier. Finally, we compare the utility of graph embeddings from different protein-folding models.

## Background

Rare diseases, while individually uncommon, collectively affect about 5% of the global population, or approximately 350 million people ^1^. With around 70% of these cases involving children the importance of early diagnosis and treatment cannot be overstated ^1^. However, the current average diagnosis time still stands at a distressing 5-7 years ^1^. Roughly 80% of these rare diseases have a genetic cause ^1^. Given the complexity of our genome, with its 3.3 billion bases and each individual carrying about 3.7 million variants, about 10.000 non-synonymous variants within the protein-coding region, pinpointing the single disease-causing variant is extremely challenging ^2^. This is further complicated by the fact that for the majority of known missense variants, a frequent cause of disease, the impact on protein function is unclear and the variants are therefore classified as variants of unknown clinical significance (VUS) ^3^. This frequently leads to inconclusive results when genetic testing is performed in a clinical setting ^4^. To better characterize missense variants and to eventually enter an age without variants of unknown clinical significance different strategies have been proposed ^5^. Ultimately experimental characterizations might be necessary to achieve this state. However, experimental approaches are resource and time consuming and consequently don’t offer promises for scalability in the near future ^6^. Therefore, different bioinformatic prioritization strategies have been proposed to narrow down the number of candidates for experimental follow ups ^7,8^. Especially machine learning models offer a promising avenue for enhancing the process of detecting and prioritizing variants ^6,9^. Pathogenicity prediction models typically include a range of genomic features, which are aggregated and delivered to a statistical or machine learning model that performs a classification or regression task ^8^. These features can include population metrics (e.g. the population allele frequency of a variant), evolutionary conservation metrics, the sequence context and epigenetic data. However, despite the progress in computational prediction strategies (e.g. AlphaFold ^10^) three-dimensional data is rarely used. Current models that use this information are either computationally expensive and bound to structure predictions from specific models (AlphaMissense ^6^) or do not use the structure itself, but information derived from this structure (SIGMA ^11^, AlphScore ^12^). Furthermore, in the case of AlphaMissense and AlphScore the models are only presented with wild type structures during training ^6,12^. While it has been previously demonstrated that variant structures predicted by AlphaFold2 don’t always agree with experimental data ^13^, models like SIGMA highlight the potential of in silico predicted variant structures ^11^. In this study, we introduce a novel machine learning approach that directly leverages information from in silico predicted protein structures of missense variants and their corresponding wild-type structures. Importantly, we used ESMFold ^14^ to predict over 60,000 protein structures to aid this process. In addition, it incorporates features derived from population genomics to construct a binary classifier for estimating the pathogenicity of missense variants. We demonstrate the practicality and value of using in silico predicted protein structures, such as those modeled by ESMFold ^14^, as an additional feature enriching machine learning approaches. This study further highlights the potential of machine learning in aiding the diagnosis of rare diseases.

## Materials

ProteinGym is a large-scale dataset published by Notin et al., that serves as benchmark for protein design models and fitness predictions, aiming to establish a basis for comparison across different studies ^15^. It contains aggregated deep mutational scanning assays and a smaller clinical dataset, derived from expert curated variant collections. Both sets are available for missense and indel variants. Additionally, it provides a benchmark board, where different pathogenicity prediction tools are ranked. For this study, we utilized the clinical substitution dataset, which contains 63,914 missense variants. The dataset consists of 31,546 variants that are classified as benign, and 32,638 variants ranked as pathogenic variants. The included variants affect 2,525 genes in total. The features included in ProteinGym are the wildtype amino acid sequence, the mutant amino acid sequence, the reference amino acid and the mutant amino acid at the place of substitution and the protein position of the substitution.

The genome aggregation database (gnomAD) includes information about 16,412,219 missense variants in its gnomAD v4 release ^16^. We extracted the population allele frequency from this release for all variants included in the clinical substitution dataset and merged them with the features already available in ProteinGym. Additionally, we further enriched the dataset by gathering the missense observed / expected ratio, obtained from the gnomAD 2.1 release ^16^. The AlphaFold database contains over 214 million predicted protein structures, as of 2024 ^17^. The structures were predicted using the AlphaFold2 model from DeepMind. We utilized the id mapping feature from UniProt ^18^ to map the proteins present in ProteinGym (identified by the NCBI protein id) to prefolded structures from the AlphaFold database. This mapping was successful for 2411 unique proteins present in ProteinGym, which translated to 59,525 out of 63,914 variants. Experiments containing the AlphaFold2 structures were therefore limited to this subset of ProteinGym.

The Zoonomia ^19^ project was started to identify conserved genomic regions, across multiple species. In their recent iteration, the international collaboration group aligned genomes for 240 species and annotated genomic regions with a score (the phyloP score), which quantifies the evolutionary conservation on a nucleotide level. Using the previously introduced id mapping feature from UniProt, we extracted the genomic correlates of the proteins present in ProteinGym. This enabled us to map the amino acid positions to three-nucleotide aggregated phyloP scores and therefore utilize phyloP scores as an additional feature in the pathogenicity prediction. This mapping was successful for 56,732 out of 63,914 variants. Experiments containing these phyloP values were therefore limited to this subset of ProteinGym.

## Methods

The presented workflow started with a protein language model. ESMFold was used to generate in silico predictions for the protein structures of the wild types and structures of the variants contained in the clinical substitution dataset from ProteinGym. ESMFold was selected, because it provides a much faster folding time and is more portable when compared to AlphaFold2, although it comes with slightly less accurate predictions. The generated structures were used to train graph autoencoders ^20^ in order to generate structural embeddings for both, the wild types and their corresponding variants. These structural embeddings were then combined with population metrics and a subset of features contained in ProteinGym. The combined features were finally used to train a XGBoost ^21^ model on a binary classification task to predict the pathogenicity of missense single nucleotide variants. The complete workflow is detailed in figure 1.

**Figure 1:**
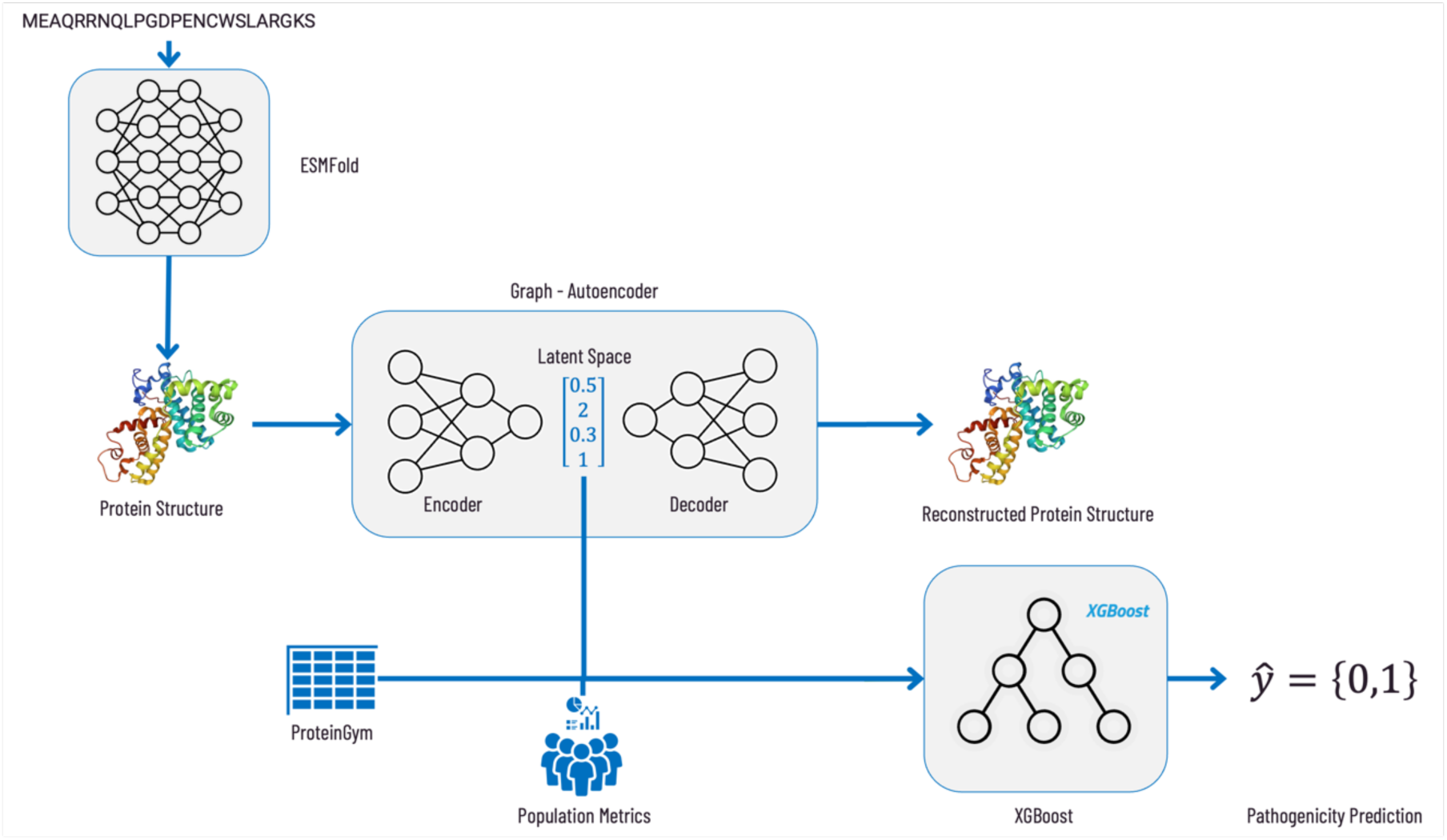
Schematic illustration of the machine learning approach workflow. We predicted protein structures for 63,914 missense single nucleotide variants and their corresponding wild types using ESMFold. These structures were then converted into numerical representations, so called embeddings, via a graph autoencoder. Finally, an XGBoost classifier was trained using these embeddings along with features from gnomAD and ProteinGym.

### Protein structure prediction with ESMFold

To generate the protein structures, we employed the ESMFold model (3B) from Hugging Face ^22^, which was utilized to predict the protein structures for both the variants and corresponding wild types within the ProteinGym dataset. This process was conducted on multiple Virtual Machines (VMs) within the Azure Machine Learning environment, each equipped with an A100 GPU.

Given the O(n^3) complexity of the ESMFold model, attempting to run inference on larger amino sequences resulted in memory overflow issues ^14^. To circumvent this, we divided longer sequences into smaller subsequences, running the inference on each subsequence individually.

The individual predictions were subsequently stitched together in a post-processing step using the NumPy library. The resulting protein structures were stored as Protein Data Bank (PDB) ^23^ files for subsequent processing and analysis in our study. This strategy allowed us to efficiently manage computational resources while generating a comprehensive set of protein structure predictions for our machine learning approach.

### Preparation of Protein Graph Datasets

We transformed the predicted protein structures from ESMFold into graph datasets represented as PyTorch Geometric Objects ^24^ using Graphein ^25^. In more detail two distinct graph datasets were generated to extract structural embeddings of varying scopes as shown figure 2:

1. An atomic-scoped dataset, where individual atoms are represented by each node, and covalent edges are based on atomic distances. The node features included the 3D coordinates and a one-hot encoding of the atom. The min-max scaled atomic distances were also included as edge features.
2. A residue-scoped dataset, where each node represents a residue in turn depicted by the alpha carbon and edges signify various interactions – distance based, aromatic, hydrogen bond, hydrophobic, aromatic Sulphur, disulfide, cation pi and peptide bonds. Node features included the 3D coordinates, the one-hot encodings of the amino acid and details about the residués presence of a hydrogen bond acceptor/donor.

**Figure 2:**
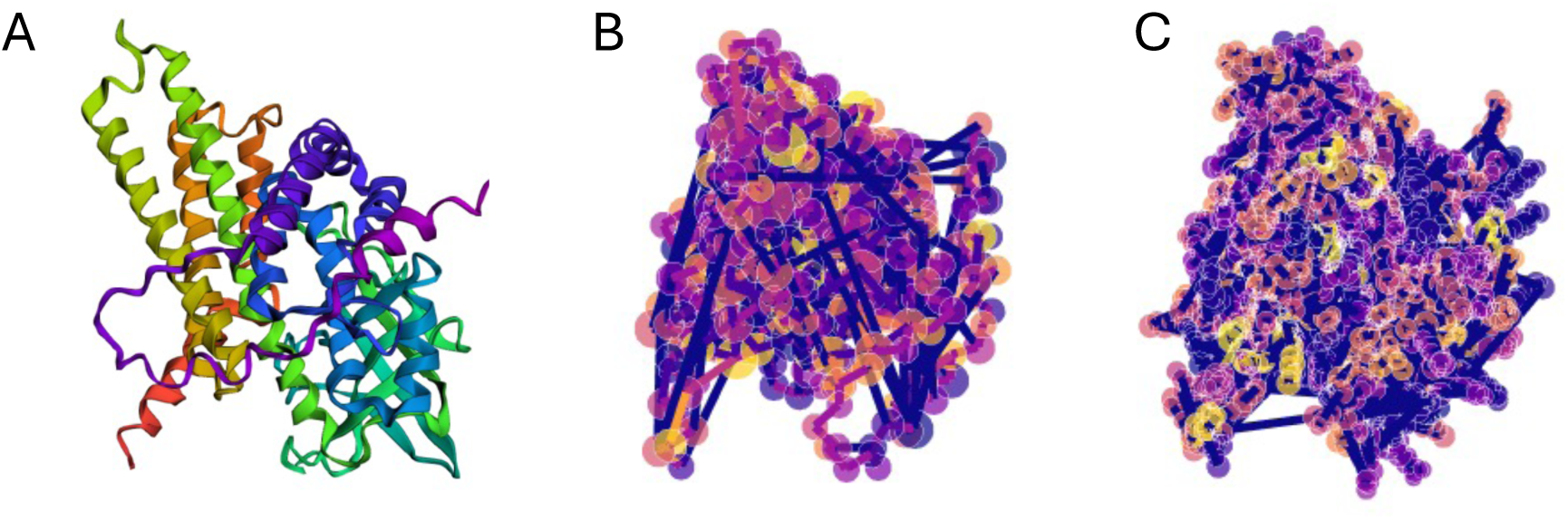
Protein Structure of A113D (A) consisting of 421 amino acids and its conversion in a residue-scoped graph representation (B) with 421 nodes and 1136 edges as well its conversion in an atomic-scoped graph representation (C) with 3274 nodes and 3338 edges. Conversion was carried out using Graphein.

**Figure 3:**
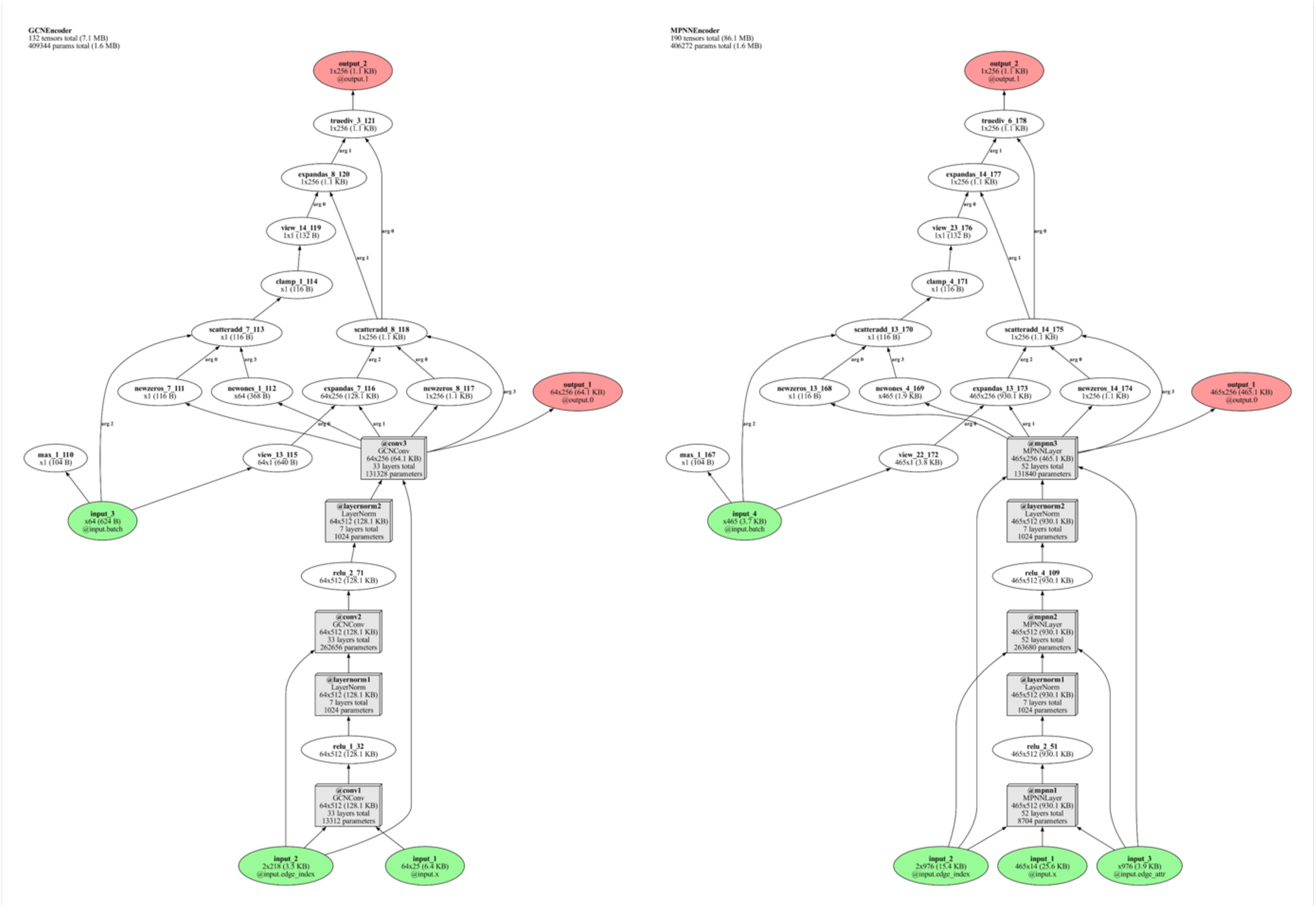
Schematic reprensentation of the GCEncoder and the MPNNEncoder, created using torchlens *^2C^*.

Both graph datasets, containing wild type and variant structures, were divided into subsets for training, validation, and testing of the graph autoencoders (GAEs). The division was conducted at the level of individual structures in a 70% (training), 20% (validation), and 10% (test) ratio.

### Structural Embeddings with Graph Autoencoders

1. Two graph autoencoders were designed, one for each graph dataset. Both architectures were implemented using PyTorch Geometric^24^ each composed of distinct custom encoders and the same inner product decoder^20^. The graph convolutional encoder (GCEncoder), designed to handle residue-scoped graphs, comprised of graph convolutional network (GCN) layers, each followed by a layer normalization applied per graph. A rectified linear unit (ReLU) activation function was applied after each GCN layer, excluding the final one. The node embeddings from the final GCN layer were pooled using a global mean pooling operation to obtain a graph-level embedding. To create graph embeddings of dimension 128 two GCN Layers, for embeddings of dimension 256 three GCN Layers were used.
2. To accommodate atomic-scoped graphs and incorporate edge features, the simple GCN layers in the GCEncoder were replaced with message passing neural network (MPNN) layers, resulting in a new encoder, the MPNNEncoder. The MPNN layers consisted of a GCN for node features and a linear layer for edge features, facilitating the transformation and integration of both node and edge information, which enhanced the model’s ability to capture more intricate graph structures. Like the GCEncoder, for each MPNN layer a ReLU activation was applied followed by layer normalization, except for the final one. The node embeddings from the final MPNN layer were pooled to achieve a graph-level embedding. To create graph embeddings of dimension 128 two MPNN Layers, for embeddings of dimension 256 three MPNN Layers were used.

Irrespective of the encoder type, an inner product decoder was utilized to decode the node embeddings, or latent variables, into edge probabilities and a probabilistic dense adjacency matrix.

Both graph autoencoders (GAE) were trained to minimize the binary cross-entropy loss for positive edges and negative sampled edges. Thus, the reconstruction loss was calculated as the sum of the losses for positive and negative edges. The GAEs were trained over a maximum of 20 epochs using a batch size of 32 and a learning rate of 0.005 with the Adam optimizer. Early stopping was implemented and evaluated epoch wise. The decision to apply early stopping was based on validation accuracy with a patience latency of 3 epochs without an increase in validation accuracy. The training, validation, and test data were loaded using PyTorch’s DataLoader^24^, which provided the data in mini batches during training. After training, the models were tested using the test data, and the Area Under Curve (AUC) and Average Precision (AP) scores were reported. Upon completion of training, the entire graph datasets, representing all predicted protein structures (wildtype and variant structures), were processed through the trained GAEs. This step enabled us to extract numerical embeddings for both the variant and wildtype structures across the entire datasets. These embeddings provide a condensed yet comprehensive representation capturing both local and global information of the protein structures, serving as a crucial input for subsequent analysis and training of a XGBoost Classifier^21^.

### Setting up a five-fold cross validation

The total dataset was split into five folds, which were eventually used to set up a five-fold cross validation. To prevent intergenic data leakage, we ensured that variants of genes present in one fold did not occur in another fold, which could have resulted in the problematic situation that variants from the same gene would end up in training, validation and test sets. This step was critical to ensure that our model’s performance evaluation was accurate and not influenced by any overlapping data between the training, validation and testing phases. While the five folds were equally sized in terms of genes per fold, genes contained in the ProteinGym clinical substitution dataset don’t contain the same number of variants and as previously mentioned benign and pathogenic variants are not equally distributed in the ProteinGym either. This resulted in slight class imbalanced folds. To balance out this imbalance, benign and pathogenic variants were randomly resampled from each fold, to bring all folds to the same variant size and an equal class distribution. The fold, which would eventually be used as current test set was not resampled, to avoid distortion of performance metrics which could result from duplicates in the test set.

For the experiments containing AlphaFold2 structures or phyloP values, we created separate folds, to ensure that the proteins and variants lost in the id mapping process were not spread unevenly across the different folds.

### Implementation and training of a XGBoost classifier

To predict the pathogenicity of missense single nucleotide variants, we adopted an XGBoost Classifier ^21^ using the XGBoost library, a gradient boosting framework renowned for its predictive accuracy and computational efficiency. For the training, validation and evaluation we used a five-fold cross validation. Followingly an XGBoost model was trained and evaluated five times using the prepared folds. For each training setup an individual data split was performed. The training set contained three folds and the validation as well as the test set contained one fold. The hyperparameters of the classifier were optimized using Optuna ^27^, a hyperparameter optimization framework. The Optuna ^27^ hyperparameter optimization was conducted over 100 trials to determine the optimal set of hyperparameters for the XGBoost^21^ model. The performance metric for the optimization was the accuracy of the model, deployed on the hold out validation set. This optimization was performed individually for each fold-split. Followingly, the five sets of tuned hyperparameters were averaged and used to train a meta-optimized model on the fivefold splits, which was used to evaluate the final performance of the classification model. This procedure was performed for different feature combinations, each with consistent fold-splits. In the first experiment setup the dataset consisted of various features including encoded amino acid references, encoded amino acid alternatives, amino acid position, the structural embeddings of both the variant and wild-type proteins as well as the cosine distance of these structural embeddings. The target variable was the pathogenicity of the variants, encoded into numerical form. We repeated this experiment with the same set up, with the exception of the inclusion of the structural embeddings and the cosine-distance of these structural embeddings, as features, to examine the added effectiveness of three-dimensional-information on a XGBoost ^21^ classifier. The final performance of each model was evaluated using the test datasets. The performance of each model on the test datasets was averaged across the folds and reported in the form of the average area under the curve of the receiver operating curve (AUROC) ^28^. Models, their hyperparameters, and their performance metrics were logged and stored using MLflow ^29^, a platform for managing the machine learning lifecycle. This approach allowed us to efficiently manage, track, and evaluate the performance of our machine learning models. The averaged hyperparameters for each experiment can be found in the supplementary information, attached to this article.

### Computation of Shapley Values

To understand the feature importance and overall model impact per feature in our XGBoost Classifiers we computed the SHAP values. The SHAP values were calculated for each fold utilizing the individual test set in the fivefold cross-validation process. These values were then stacked across all folds to assess the overall feature importance across the whole dataset. By default, the SHAP values are computed for each input feature, which in our case corresponded to each element in our structural embeddings.

Given the additive nature of the SHAP values, we wanted to consider the structural embeddings as a whole. To achieve this, we summed the SHAP values over each structural embedding space. This allowed us to obtain a single SHAP value per structure embedding, providing a comprehensive understanding of the importance of the entire structural embedding in the model. These steps allowed us to create feature importance plots, which visually represented the significance and total impact of each feature in the model.

## Results

### Pathogenicity Prediction

To access the value of in silico predicted three-dimensional protein structures for pathogenicity prediction of missense variants, we trained multiple XGBoost ^21^ classifiers. Each model was presented with population metrics and, aside from one exception of one (the minimal model), was additionally enriched with structural graph embeddings of different abstraction levels. The performance of all models was subsequently evaluated using the AUROC metric. The individual plotted ROC-Curves for all classifiers can be seen in figure 4. The AUROC of the classifiers, that were additionally supplemented with the structural graph embeddings showed a slight, but consistent tendency towards an increase in performance, as demonstrated by the comparison of the average AUROCs. The classifier, that was trained and evaluated without the structural graph embeddings (the minimal model) showed the following AUROC values across the individual folds [fold1=0.92, fold2=0.92, fold3=0.90, fold4=0.90, fold5=0.89] and a mean AUROC value of 0.906 (standard deviation = 0.011). The overall best performing classifier was the classifier additionally supplemented with both the graph embeddings on atomic level of abstraction (embedding size = 128) and the graph embeddings on residue level of abstraction (embedding size = 128). It reached the following AUROC values across the individual folds [fold1=0.93, fold2=0.92, fold3=0.91, fold4=0.93, fold5=0.93] and a mean AUROC value of 0.924 (standard deviation = 0.009). However, the difference to the second-best preforming model, the classifier supplemented with graph embeddings on the residue level of abstraction (embedding size = 128), was marginal. This second-best performing classifier reached the following AUROC values across the individual folds [fold1=0.93, fold2=0.93, fold3=0.90, fold4=0.92, fold5=0.92] and a mean AUROC value of 0.922 (standard deviation = 0.010).

**Figure 4:**
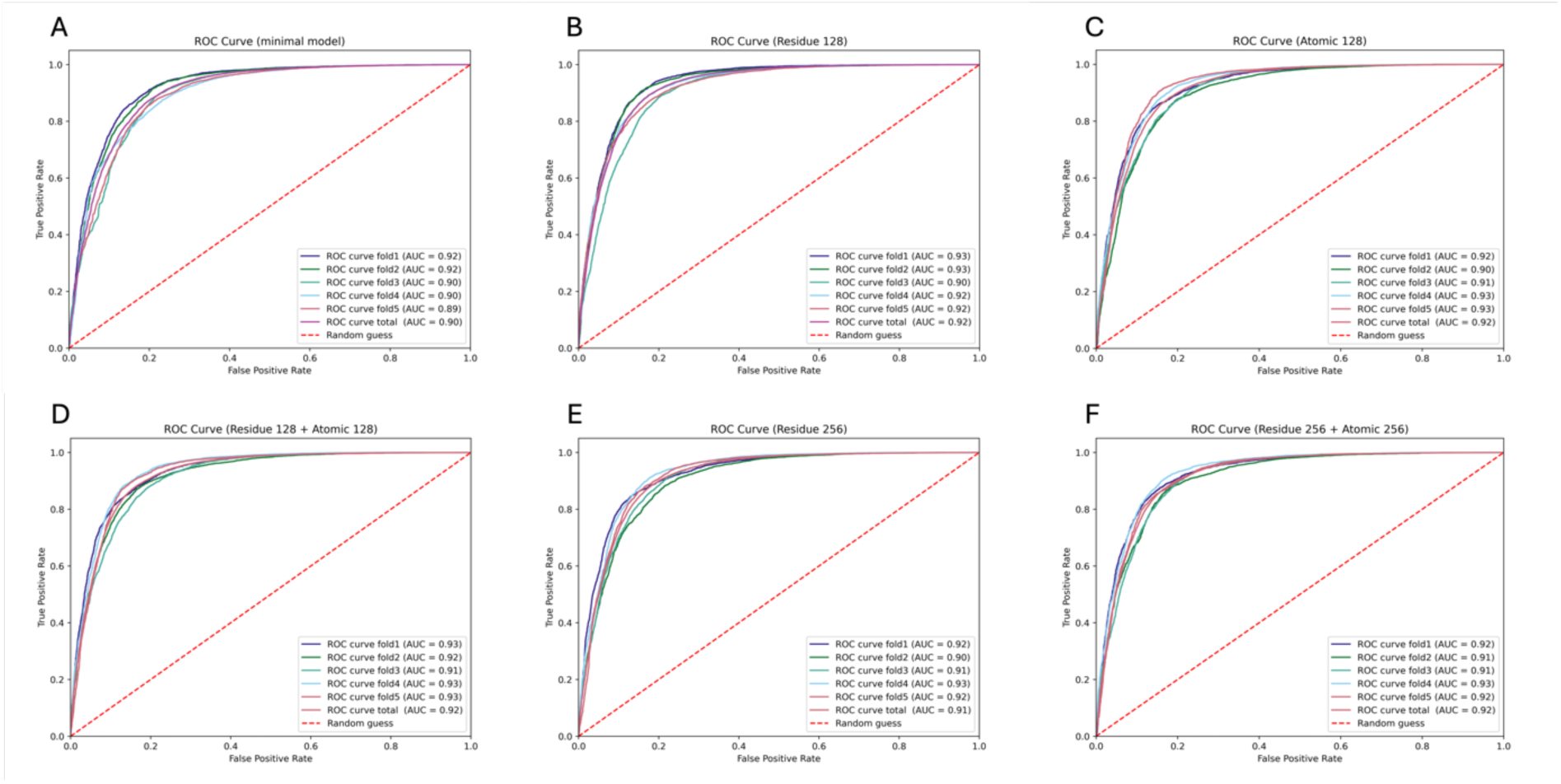
Side-by-Side comparison of the performance of the different classifier models. (A) Classifier trained without the graph embeddings. (B) Classifier trained with graph embeddings on the residue level of abstraction (embedding size 128). (C) Classifier trained with graph embeddings on the atomic level of abstraction (embedding size 128). (D) Classifier trained with graph embeddings both on residue level of abstraction (embedding size 128) and atomic level of abstraction (embedding size 128). (E) Classifier trained with graph embeddings on residue level of abstraction (embedding size 256). (F) Classifier trained with graph embeddings both on residue level of abstraction (embedding size 256) and atomic level of abstraction (embedding size 256). The ROC-Curve of the Model trained on the graph-embeddings on atomic level of abstraction with the larger embeddings size of 256 can be found in supplementary figure 1.

### Exploration of different embedding sizes

As previously stated, and depicted in figure 4, next to different level of abstractions of the protein graphs, we additionally analyzed whether the embedding size has an impact on the performance of utilized classifiers. In general, we observed a small yet consistent difference in the side-by side comparison between classifiers on the same abstraction level but different embedding sizes. Overall, the classifiers trained on smaller embedding size showed an equal or slightly higher mean AUROC value, when compared to their counterpart trained on larger embeddings sizes. This was consistently observed in the residue-by-residue comparison (mean AUROC residue 128 = 0.923 standard deviation = 0.010 & mean AUROC residue 256 = 0.916; standard deviation = 0.009), the atomic-by-atomic comparison (mean AUROC atomic 128 = 0.917; standard deviation = 0.010 & mean AUROC atomic 256 = 0.917; standard deviation = 0.005) and also in the mixed-by-mixed comparison (mean AUROC residue 128/ atomic 128 = 0.924; standard deviation = 0.009 & mean AUROC residue 256 / atomic 256 = 0.918; standard deviation = 0.008). No model trained on the larger embedding size of 256 outperformed their 128 embedding size counterparts. An overall comparison in the form of a bar plot can be seen in figure 5.

**Figure 5:**
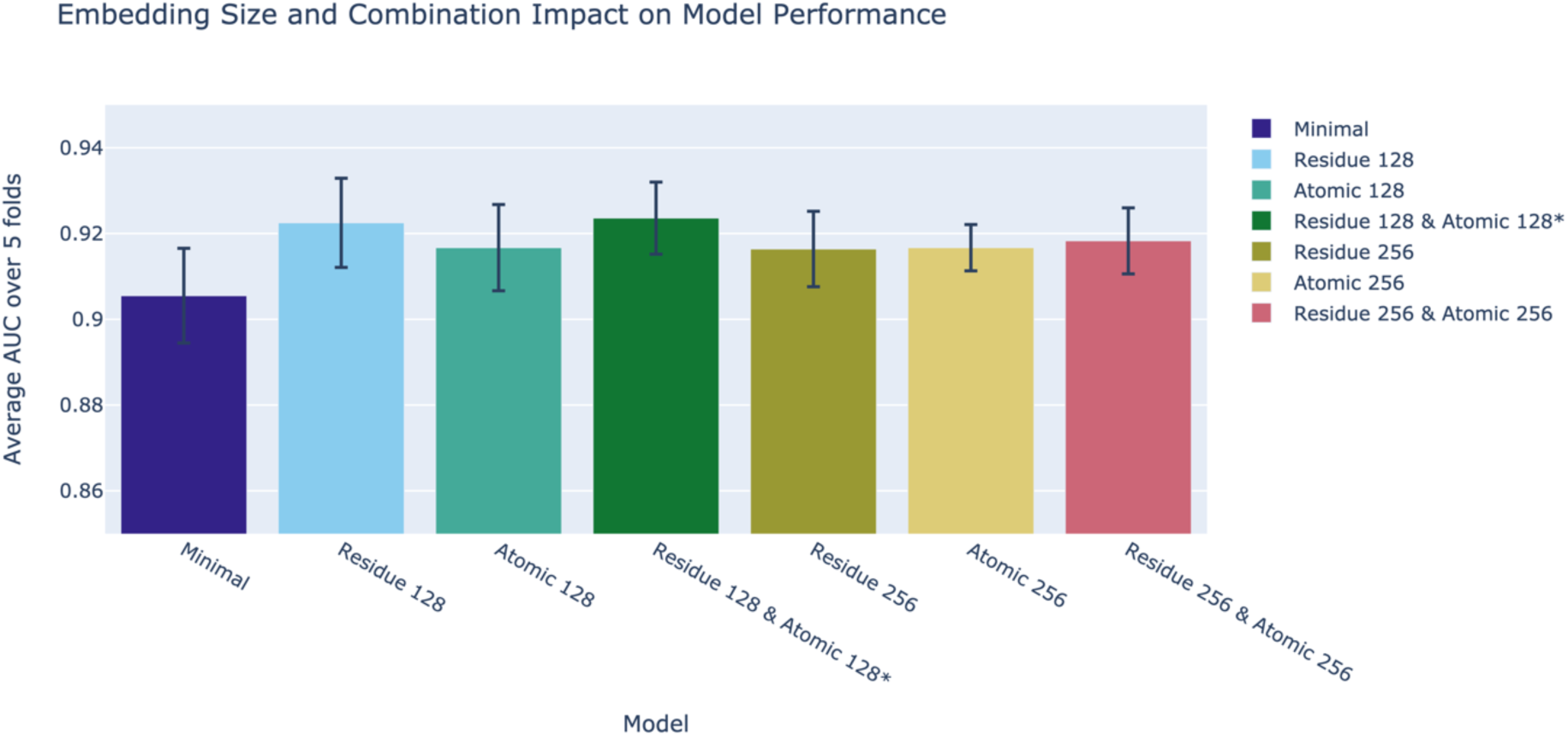
Bar plot highlighting the AUROC differences on the different combinations of level of abstractions and embedding sizes. Overall, the experiments utilizing smaller embeddings showed slightly higher AUROC values. The best performing classifier utilized embeddings from both levels of abstraction, although the difference was minimal. Next to the AUROC values, the standard deviation is depicted

### Feature Importance

To further explore the relevance of graph embeddings for the classification task, we utilized the individually fold wise trained XGBoost classifiers at the residue level of abstraction (embedding size = 128) to predict the SHAP values for the hold out test data. We aggregated these SHAP values for an overall evaluation which can be seen in figure 6. As visualized in figure 6, the allele frequency is the most influential feature for pathogenicity prediction, which is then followed by the structural embeddings of the wild-type and mutant structures.

**Figure 6:**
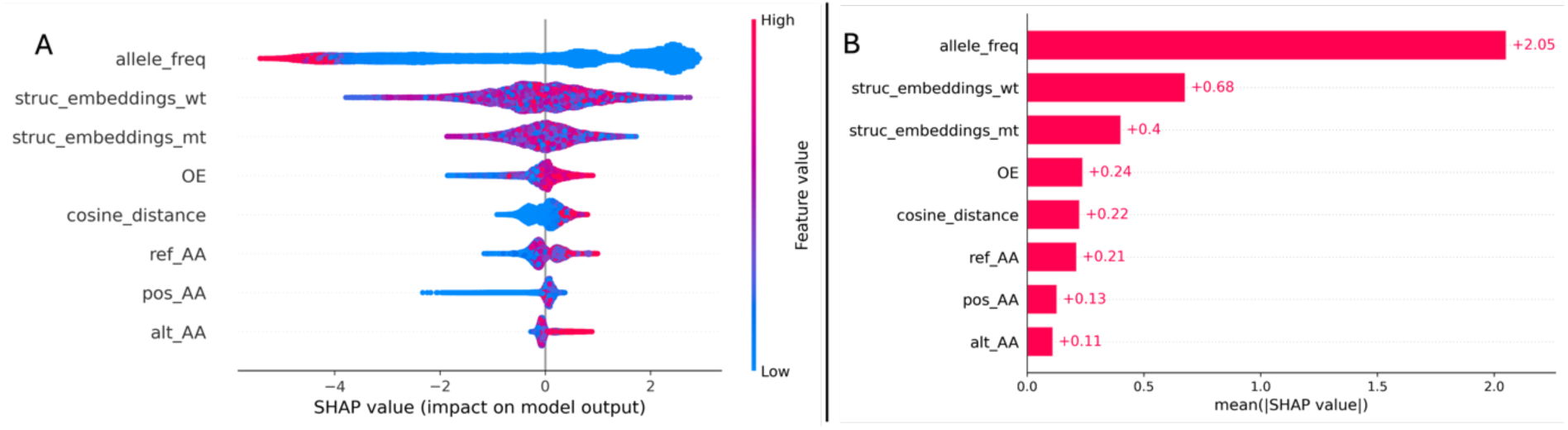
SHAP values displayed as a bee swarm plot (A), highlighting the individual SHAP values. Additionally, the aggregated SHAP values are displayed as bar plot (B). The allele frequency is the undisputed most important feature. The graph embeddings are of noticeable importance. The wild type structures are ranked as more important to the classification task, compared to the variant structures.

### Comparison to previous scores

ProteinGym is a standardized dataset which can be used to evaluate and compare different pathogenicity predictors. However, for the clinical substitutions’ dataset no predefined cross validation folds or train-test splits are available. Additionally, different models evaluated on ProteinGym are often trained on additional data and data leakage is not always considered, leading to a leaderboard which overestimates the performance of some classifiers. A comparison of our best performing XGBoost to other models listed on the ProteinGym leaderboard can be seen in figure 7.

**Figure 7:**
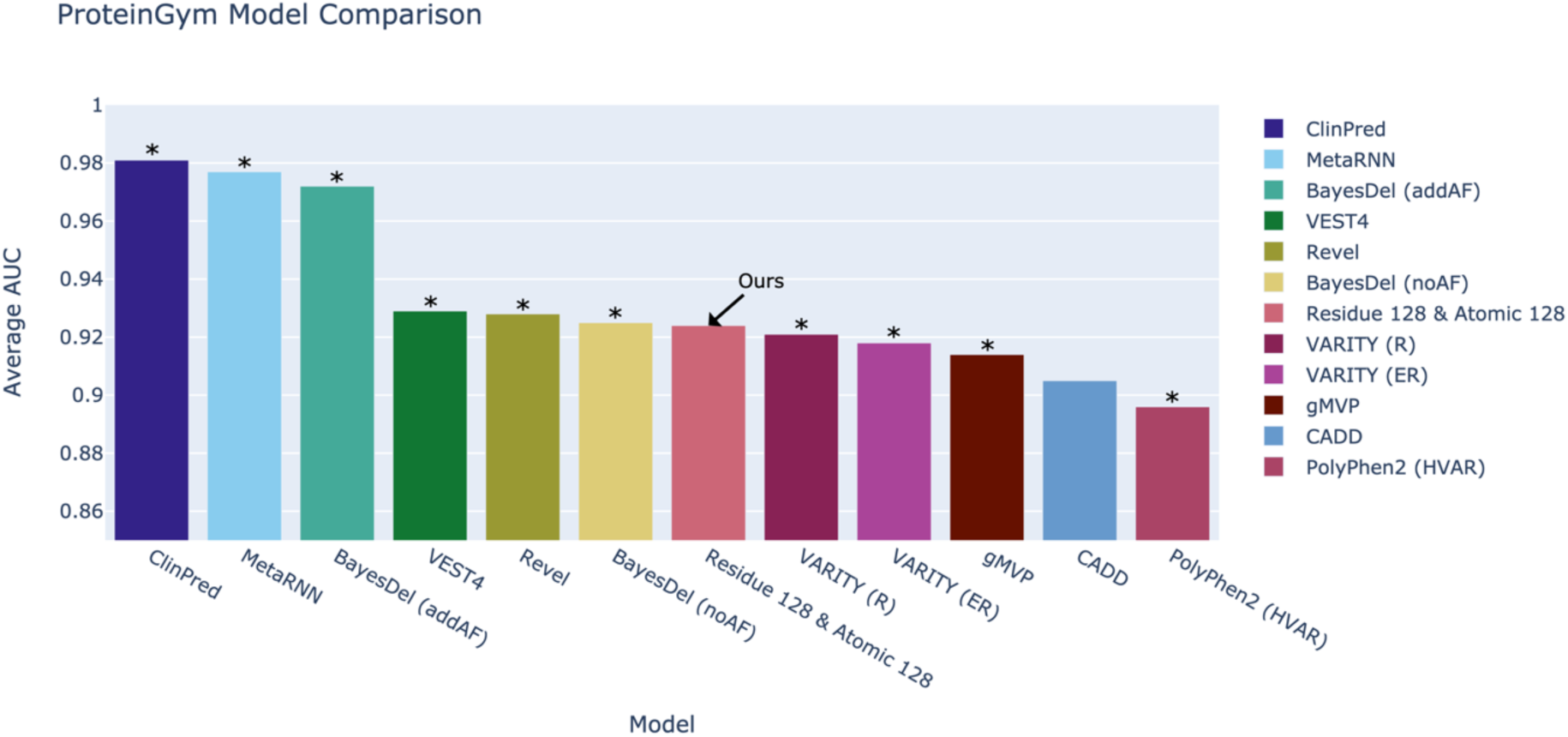
ProteinGym leaderboard as depicted on the ProteinGym website ^7,8,30–36^. Since no standard procedure is defined for training and evaluation on the clinical substitution dataset, and some of the models predate the introduction of ProteinGym, most models are trained on additional data. Models which were trained on larger datasets e.g. the entirety of ClinVar which has a large overlap with the clinical substitution dataset leading to differing magnitudes of data leakage are marked with an asterisk (*).

### Comparing graph embeddings from different protein folding models

While most of the analysis in this manuscript were focused on in silico structures predicted by ESMFold, we hypothesized that the presented workflow is agnostic towards the source of protein structures. To put this hypothesis to the test we obtained wildtype structures predicted by AlphaFold2 for the proteins present in ProteinGym. We embedded these Alphafold2 structures utilizing the autoencoder model, trained on ESMFold data. Utilizing these structures we repeated the experiments (with residue level of abstraction and embedding size of 128), with the difference that only ESMFold or AlphaFold2 wildtype structures were used to augment the other non-structure related features. The classifier presented with ESMFold wildtype structures reached a mean AUROC of 0.918 (standard deviation = 0.012), while the classifier supplemented with AlphaFol2 wildtype structures reached a mean AUROC of 0.921 (standard deviation = 0.007) The resulting ROC-Curves can be seen in figure 8.

**Figure 8:**
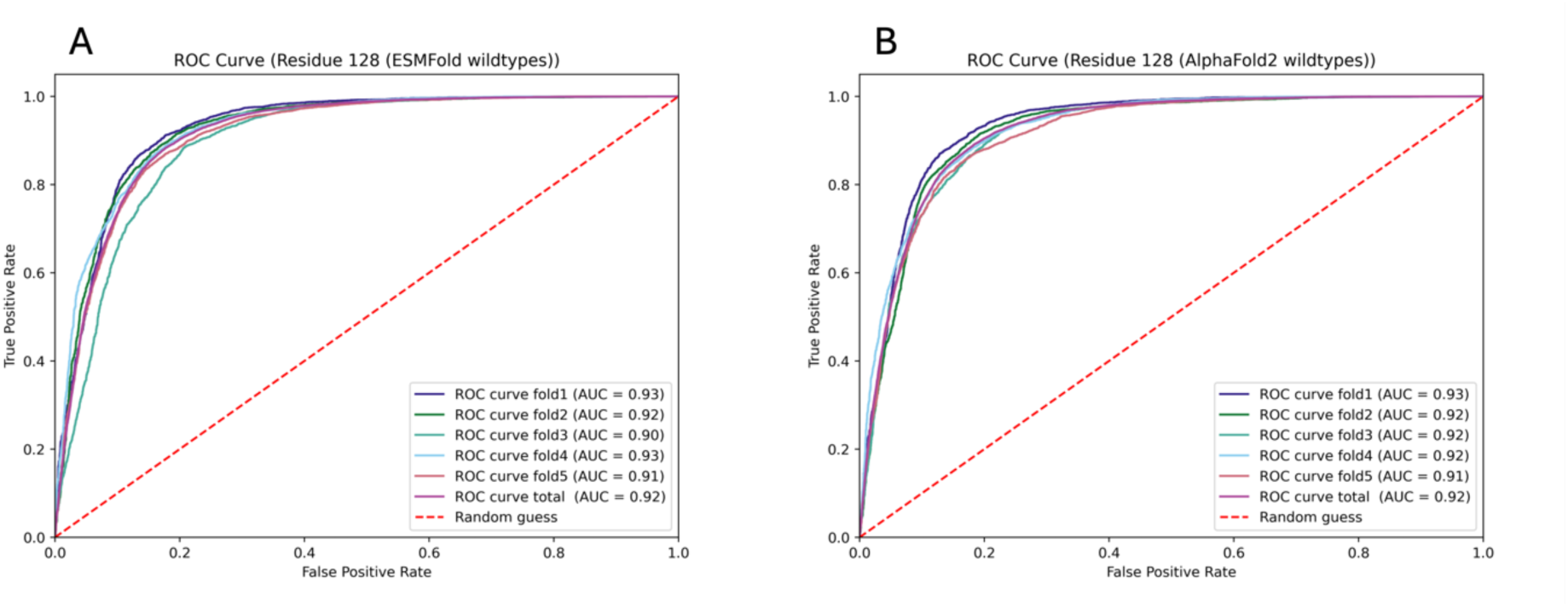
(A) ROC-Curve for XGBoost classifier trained on data from ProteinGym, gnomAD and wildtype structures predicted by ESMFold. (B) ROC-Curve for XGBoost classifier trained on data from ProteinGym, gnomAD and wildtype structures by AlphaFold2.

### Extending the feature space by evolutionary conservation scores

To estimate how additional features might influence the importance of structural embeddings, we additionally supplied the classifier with a measurement of evolutionary conservation. To be more precise we added the latest phyloP scores from the Zoonomia ^19^ project as an additional feature for pathogenicity prediction. Using the extended feature space, we repeated the initial experiments for both the minimal model and the residue level of abstraction with embedding size 128. The new minimal model reached an AUROC value of 0.916 (standard deviation = 0.011) while the phyloP extended residue level of abstraction model reached an AUROC value of 0.932 (standard deviation = 0.009). The respective ROC-Curves can be seen in figure 9. To analyze whether the addition of the phyloP scores reduced the impact of the structural embeddings on the model, we additionally obtained SHAP values for this new model. The SHAP plots can be seen in figure 10.

**Figure 9:**
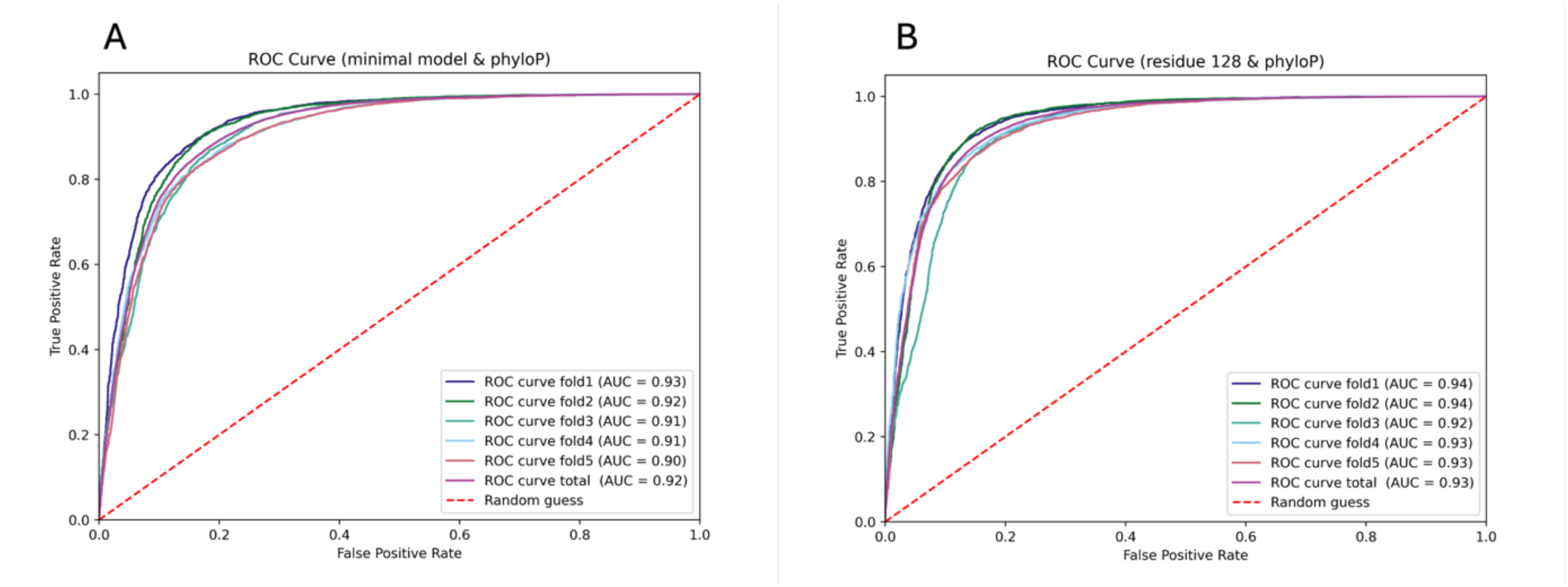
(A) ROC-Curve of the new minimal model, which was trained using the extended feature space by pyhloP values from the Zoonomia project. (B) ROC-Curve of the classifier using the extended feature space by phyloP values and supplemented with structural embeddings for variant and wildtype structures, presented on residue level of abstraction, with the embedding size of 128.

**Figure 10:**
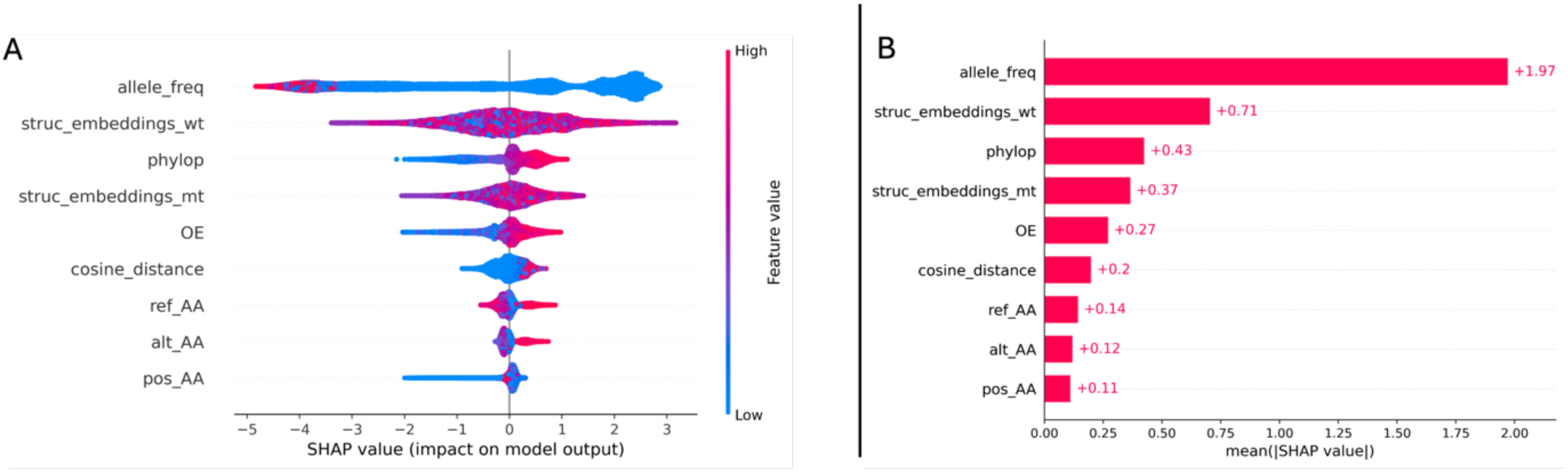
SHAP values displayed as a bee swam plot (A), highlighting the individual SHAP values. Additionally, the aggregated SHAP values are displayed as a bar plot (B). Compared to figure 6, the newly added phyloP score overtakes the variant structural embeddings, in terms on absolute influence of the model.

## Discussion

### Differentiating to existing models

The demonstrated workflow presents a way to include the three-dimensional structure of proteins in pathogenicity classification tasks. In principle this agnostic workflow is capable of processing experimentally determined and in silico predicted structures. Models like AlphaFold2 ^10^ and ESMFold ^14^ have made in silico predicted protein structures abundantly available, however these structures have only been partially used in variant effect predictors previously. Previous structural aware models like those presented by Schmidt et al. and Zhao et al. take structural information from in silico predicted protein structures from AlphaFold2 into account, however they rely on an engineered feature extraction process in which biochemical and network features are extracted and then used to train a classifier model^11,12^. The presented approach differentiates itself from those previous workflows by replacing the manual feature engineering process with graph embeddings generated by graph autoencoders, which is compatible with in silico predicted structures from arbitrary computational modelling approaches and real-world structures.

### Exploring protein graph embeddings at different levels of abstraction

We examined whether the level of abstraction used to transform in silico generated protein structures to protein graphs, has different effects on a classifier used for pathogenicity prediction. While the overall best performing model, in our initial experiments, was a model trained on a combination of atomic level structural embeddings and residue level structural embeddings, the improvement was minimal, when compared to a model trained solely on residue level structural embeddings. For the smaller embedding size of 128, models trained only on the atomic level structural embeddings, were outperformed by their residue counterparts, although again the differences between the model’s performance were small. This could have different explanations. Although it has been demonstrated that variant structures from protein-folding models contain useful information that can be leveraged in downstream tasks ^11,12^, protein-folding models still fall a little bit short when evaluated on the atomic scope ^37–40^. We therefore hypothesize that the most likely explanation in our case would be that embeddings generated from structures generated by the ESMFold model on the atomic scope, do not accurately capture actual atomic constellations. This effect is probably further amplified in variant structures on the atomic scope, because as introduced they already provide difficulties for current models. Future work is needed in which this is explored in more detail, possibly even in a comparison between in silico and real-world structures.

### Comparing protein graph embeddings from different sources

Due to computational limitations we opted to use the 3 billion parameter ESMFold model, available on HuggingFace^22^. Because of the faster inference and smaller computational requirements, when compared to AlphaFold2, we were able to predict the wildtype and variant structures for the whole ProteinGym clinical substitution dataset. This however is a possible entry point for a performance ceiling effect, since for ESMFold it has been demonstrated that an increase in parameter size results in more accurate predictions. Additionally, since its release, ESMFold has been described as slightly less accurate compared to AlphaFold2 ^14^. This reduced accuracy might be inherited by the presented classifiers and therefore should be kept in mind as potential limitation. As previously introduced, we hypothesize this workflow to be somewhat source agnostic. We provide evidence for this protein structure agnosticism with our AlphaFold2 experiment, by demonstrating that a classifier trained with AlphaFold2 graph embeddings (generated using an autoencoder trained purely on ESMFold data), achieves similar performances when compared to a classifier trained on ESMFold graph embeddings (generated using an autoencoder trained purely on ESMFold data). Because of limited computational resources, we did not have the capacity to additionally fold all variant structures utilizing AlphaFold2 or even AlphaFold3, ^41^ which would be an interesting comparison to be made in a future project. We justify this short-cut by information obtained from figure 6. While both types of structural embeddings (wildtype and variant) influence the model’s performance, the wildtype structures seem to have a larger impact. This larger impact was observable across all our experiments. Therefore, we estimated that the wildtype structures alone can be used to obtain evidence for generalizability across different structures from different sources. In future work, this idea needs to be examined by performing experiments containing wildtype and variant structures from different sources.

### Protein structures as a utility in pathogenicity prediction classifiers

In a final experiment we examined how the influence of the structural embeddings behaves, once (in addition to the general allele frequency from the population) a strong feature such as a measurement of evolutionary conservation is added to the classifier. We demonstrate that the impact of the wildtype structures remains largely unchanged, as demonstrated by the calculated SHAP-values. While the phyloP score surpasses the variant structural embeddings in their impact as presented in figure 10, the absolute value changes only slightly, when compared to the value present in figure 6. This is further supported by the fact, that the overall best performing model in this study is a classifier which combines all presented features, which might highlight that the structural embeddings contain some kind of information, which is not adequately captured by the population frequencies and information about evolutionary conservation.

### Further Directions G Limitations

The presented work mainly serves as a machine learning workflow with the aim to further establish the integration of in silico predicted structures for downstream tasks like variant prioritization. For the latter, further features should be added, to develop a more complete pathogenicity prediction workflow. As a possible next step in addition to the evolutionary conservation we tried to explore post translational modifications as a possible addition. For this we obtained residue modifications through the id mapping approach from uniport. Due to the fact that there are only 335 out of 63,614 variants with affected residues that are modified following translation within the total clinical substitution dataset of ProteinGym, we concluded to postpone this feature for future versions of the outlined workflow.

Additionally, in subsequent work this workflow needs to be evaluated on datasets presented in the recent ClinGen recommendations paper ^42^, which focused on the calibration of missense prioritization scores. Finally, in principle the presented workflow is not restricted to missense or single nucleotide variants and can be applied to insertions, deletions, frameshift mutations, and multi-nucleotide variants in future studies.

### Summary

We developed a machine learning workflow that can generate graph embeddings from protein structures, which can be used in downstream tasks. We explored the utility of these embeddings in the application case of pathogenicity prediction of missense variants. We explored different levels of abstractions of these graph embeddings, compared in silico structures from different sources and evaluated the impact of combining additional features with these graph embeddings, highlighting the additional value of these graph embeddings in different experiments, by demonstrating that graph embeddings from in silico structures provide information that might be not encapsulated by other features such as the allele frequency and evolutionary conservation.

## Abbreviations

AUROC: Area under the receiving operating characteristic curve
GAE: Graph Autoencoder
GCN: Graph Convolutional Network
GCEncoder: Graph Convolutional Encoder
MPNN: Message Passing Neural Network
MPNNEncoder: Message Passing Neural Network Encoder
ROC-Curve: Receiver Operating Characteristic Curve

## Ethics approval and consent to participate

Not applicable.

## Consent for publication

Not applicable.

## Availability of data and materials

Upon publication, the code and sample data to reproduce the workflow will be available in the following GitHub repository: https://github.com/IHGGM-Aachen/genoseer.

## Authors contribution

MD and JK conceived the idea. MD and JK implemented the machine learning workflow. MD, MB and JK prepared and aggregated the different datasets. IK, ME, MB and JK performed validation and provided supervision. All authors were involved in reviewing and writing of the manuscript and gave their consent for the submission of the final version.

## References

1. The Lancet Global Health, null. The landscape for rare diseases in 2024. Lancet Glob. Health 12, e341 (2024).

2. Auton, A. et al. A global reference for human genetic variation. Nature 526, 68–74 (2015).

3. Richards, S. et al. Standards and Guidelines for the Interpretation of Sequence Variants: A Joint Consensus Recommendation of the American College of Medical Genetics and Genomics and the Association for Molecular Pathology. Genet. Med. Off. J. Am. Coll. Med. Genet. 17, 405 (2015).

4. Sessa, G., Ehlén, Å., Nicolai, C. von & Carreira, A. Missense Variants of Uncertain Significance: A Powerful Genetic Tool for Function Discovery with Clinical Implications. Cancers 13, 3719 (2021).

5. Fowler, D. M. & Rehm, H. L. Will variants of uncertain significance still exist in 2030? Am. J. Hum. Genet. 111, 5–10 (2024).

6. Cheng, J. et al. Accurate proteome-wide missense variant effect prediction with AlphaMissense. Science 381, eadg7492 (2023).

7. Ioannidis, N. M. et al. REVEL: An Ensemble Method for Predicting the Pathogenicity of Rare Missense Variants. Am. J. Hum. Genet. 99, 877 (2016).

8. Rentzsch, P., Witten, D., Cooper, G. M., Shendure, J. & Kircher, M. CADD: predicting the deleteriousness of variants throughout the human genome. Nucleic Acids Res. 47, D886–D894 (2019).

9. Schubach, M., Maass, T., Nazaretyan, L., Röner, S. & Kircher, M. CADD v1.7: using protein language models, regulatory CNNs and other nucleotide-level scores to improve genome-wide variant predictions. Nucleic Acids Res. 52, D1143–D1154 (2024).

10. Jumper, J. et al. Highly accurate protein structure prediction with AlphaFold. Nature 596, 583–589 (2021).

11. Zhao, H. et al. SIGMA leverages protein structural information to predict the pathogenicity of missense variants. Cell Rep. Methods 4, 100687 (2024).

12. Schmidt, A. et al. Predicting the pathogenicity of missense variants using features derived from AlphaFold2. Bioinformatics 39, btad280 (2023).

13. Buel, G. R. & Walters, K. J. Can AlphaFold2 predict the impact of missense mutations on structure? Nat. Struct. Mol. Biol. 29, 1–2 (2022).

14. Lin, Z. et al. Evolutionary-scale prediction of atomic-level protein structure with a language model. Science 379, 1123–1130 (2023).

15. Notin, P. et al. ProteinGym: Large-Scale Benchmarks for Protein Fitness Prediction and Design. Adv. Neural Inf. Process. Syst. 36, 64331–64379 (2023).

16. Karczewski, K. J. et al. The mutational constraint spectrum quantified from variation in 141,456 humans. Nature 581, 434–443 (2020).

17. Varadi, M. et al. AlphaFold Protein Structure Database in 2024: providing structure coverage for over 214 million protein sequences. Nucleic Acids Res. 52, D368–D375 (2023).

18. The UniProt Consortium. UniProt: the Universal Protein Knowledgebase in 2025. Nucleic Acids Res. gkae1010 (2024) doi:10.1093/nar/gkae1010.

19. Christmas, M. J. et al. Evolutionary constraint and innovation across hundreds of placental mammals. Science 380, eabn3943 (2023).

20. Kipf, T. N. & Welling, M. Variational Graph Auto-Encoders. Preprint at 10.48550/arXiv.1611.07308 (2016).

21. Chen, T. & Guestrin, C. XGBoost: A Scalable Tree Boosting System. Preprint at 10.48550/arXiv.1603.02754 (2016).

22. Wolf, T., et al. HuggingFace’s Transformers: State-of-the-art Natural Language Processing. Preprint at 10.48550/arXiv.1910.03771 (2020).

23. Burley, S. K. et al. Protein Data Bank (PDB): The Single Global Macromolecular Structure Archive. Methods Mol. Biol. Clifton NJ 1607, 627 (2017).

24. Paszke, A., et al. PyTorch: An Imperative Style, High-Performance Deep Learning Library. Preprint at 10.48550/arXiv.1912.01703 (2019).

25. Jamasb, A. et al. Graphein - a Python Library for Geometric Deep Learning and Network Analysis on Biomolecular Structures and Interaction Networks. Adv. Neural Inf. Process. Syst. 35, 27153–27167 (2022).

26. Taylor, J. & Kriegeskorte, N. Extracting and visualizing hidden activations and computational graphs of PyTorch models with TorchLens. Sci. Rep. 13, 14375 (2023).

27. Akiba, T., Sano, S., Yanase, T., Ohta, T. & Koyama, M. Optuna: A Next-generation Hyperparameter Optimization Framework. in Proceedings of the 25th ACM SIGKDD International Conference on Knowledge Discovery & Data Mining 2623–2631 (Association for Computing Machinery, New York, NY, USA, 2019). doi:10.1145/3292500.3330701.

28. Junge, M. R. J. & Dettori, J. R. ROC Solid: Receiver Operator Characteristic (ROC) Curves as a Foundation for Better Diagnostic Tests. Glob. Spine J. 8, 424 (2018).

29. Chen, A. et al. Developments in MLflow: A System to Accelerate the Machine Learning Lifecycle. in Proceedings of the Fourth International Workshop on Data Management for End-to-End Machine Learning 1–4 (Association for Computing Machinery, New York, NY, USA, 2020). doi:10.1145/3399579.3399867.

30. Alirezaie, N., Kernohan, K. D., Hartley, T., Majewski, J. & Hocking, T. D. ClinPred: Prediction Tool to Identify Disease-Relevant Nonsynonymous Single-Nucleotide Variants. Am. J. Hum. Genet. 103, 474–483 (2018).

31. Li, C., Zhi, D., Wang, K. & Liu, X. MetaRNN: differentiating rare pathogenic and rare benign missense SNVs and InDels using deep learning. Genome Med. 14, 115 (2022).

32. Feng, B.-J. PERCH: A Unified Framework for Disease Gene Prioritization. Hum. Mutat. 38, 243–251 (2017).

33. Carter, H., Douville, C., Stenson, P. D., Cooper, D. N. & Karchin, R. Identifying Mendelian disease genes with the Variant Effect Scoring Tool. BMC Genomics 14, S3 (2013).

34. Wu, Y., Li, R., Sun, S., Weile, J. & Roth, F. P. Improved pathogenicity prediction for rare human missense variants. Am. J. Hum. Genet. 108, 1891–1906 (2021).

35. Zhang, H., Xu, M. S., Fan, X., Chung, W. K. & Shen, Y. Predicting functional effect of missense variants using graph attention neural networks. Nat. Mach. Intell. 4, 1017–1028 (2022).

36. Adzhubei, I., Jordan, D. M. & Sunyaev, S. R. Predicting Functional Effect of Human Missense Mutations Using PolyPhen-2. Curr. Protoc. Hum. Genet. Editor. Board Jonathan Haines Al 0 7, Unit7.20 (2013).

37. Marcu, Ş.-B., Tăbîrcă, S. & Tangney, M. An Overview of Alphafold’s Breakthrough. Front. Artif. Intell. 5, 875587 (2022).

38. Carugo, O. Accuracy of AlphaFold models: Comparison with short N…O contacts in atomic resolution protein crystal structures. Comput. Biol. Chem. 110, 108069 (2024).

39. Wu, T., Guo, Z. & Cheng, J. Atomic protein structure refinement using all-atom graph representations and SE(3)-equivariant graph transformer. Bioinformatics 39, btad298 (2023).

40. Dai, X., Wu, L., Yoo, S. C Liu, Q. Integrating AlphaFold and deep learning for atomistic interpretation of cryo-EM maps. Brief. Bioinform. 24, bbad405 (2023).

41. Abramson, J. et al. Accurate structure prediction of biomolecular interactions with AlphaFold 3. Nature 630, 493–500 (2024).

42. Pejaver, V. et al. Calibration of computational tools for missense variant pathogenicity classification and ClinGen recommendations for PP3/BP4 criteria. Am. J. Hum. Genet. 109, 2163– 2177 (2022).

